# Multiple neural network approaches, including use of topological data analysis, enhances classification of human induced pluripotent stem cell colonies by treatment condition

**DOI:** 10.1101/2025.01.16.633346

**Authors:** Alexander Ruys de Perez, Paul E. Anderson, Elena S. Dimitrova, Melissa L. Kemp

## Abstract

Understanding how stem cells organize to form early tissue layers remains an important open question in developmental biology. Helpful in understanding this process are biomarkers or features that signal when a significant transition or decision occurs. We show such features from the spatial layout of the cells in a colony are sufficient to train neural networks to classify stem cell colonies according to differentiation protocol treatments each colony has received. We use topological data analysis to derive input information about the cells’ positions to a four-layer feedforward neural network. We find that despite the simplicity of this approach, such a network has performance similar to the traditional image classifier ResNet. We also find that network performance may reveal the time window during which differentiation occurs across multiple conditions.

**Author summary:** Our understanding of how stem cells determine what specialized cells to differentiate into is still incomplete. One aspect of understanding this process involves identifying when key decisions about a cell’s fate occur. We explored whether by looking at the layout of the cells of the colony, we can infer knowledge regarding the eventual phenotypes the cells are differentiating towards. We train an algorithm to recognize cell type using spatial information by taking as input the number and size of holes that appear among the colony’s cells. We find this method succeeds in its classification, similar to an industry-grade image classifier.

## Introduction

Organoids and microphysiological systems generated from human induced pluripotent stem cells (hiPSCs) hold promise for developing in vitro assays that can be used for evaluating therapeutics, toxicological screening, and personalized medicine. Furthermore, the way in which stem cells organize themselves into more specialized tissues under various culture conditions provides critical insight into morphogenesis, the dynamic formation of organ systems. During these processes, guidance to individual cells comes through positional cues [1] and interactions with neighbors [2]. Thus, while morphogenesis is driven by soluble morphogen signals, the process itself can be readily observed through spatial information in the form of cell migration, clustering, and changes in densities. We ask whether this spatial information can be interpreted in a way that allows us to determine when the differentiation process is precisely occurring, and what differentiation is taking place. Ultimately, the prediction of eventual lineage specification by the simple non-invasive metrics derived from multicellular organization is desired for reliability and real-time quality control of organoids in an industrial manufacturing setting.

In responding to the challenge of tissue identification from spatial data, we design a neural network which takes as input the positional data of a colony of cells, and guesses the cell fate of the colony. Applying deep learning to these image classification problems has yielded success. Image classifying neural networks have been able to identify cell morphology [3], and have found applications in cell sorting [4] and cytopathology [5]. Especially relevant to our interest is the work of [6], which predicted the differentiation of individual primary murine hematopoeic stem and progenitor cells (HSPCs) into one of two cell fates: the granulocytic/monocytic (GM) or the megakaryocitic/erythroid (MegE). In this case deep learning could accurately predict the lineage three generations before the cells began emitting the identifying markers.

We venture from the previous work by using deep learning to make conclusions about a mass of many related cells, rather than a single individual cell. Morphology has provided information about differentiation of pluripotent stem cell aggregates. In [7], classification of embryoid bodies by morphology (cystic, bright cavity, or dark cavity) predicted which early germ layers would emerge. This result introduces the potential of a computer to make further predictions based on details not evident to human observation. While a natural approach to this machine learning problem would be to design an image classifier, we also include a network that uses topological data analysis (TDA). We hypothesize that the *persistent homology* of the colony, which effectively catalogues the gaps and holes that form between the cells, will distill the critical features that would be lost in the raw image. Persistent homology has proven to be insightful in other biological contexts. Indeed, [8] found that a support vector machine could use persistent homology, when refined into a persistence landscape function, to distinguish between open and closed conformations of the maltose-binding protein. On the level of cellular interactions, [9] found that 1-dimensional homology could serve as an effective classifier for motility phases in epithelial cells.

## Results

### Both the topological data-using network and the standard image classifier succeed in classifying images

In order to investigate the potential utility of TDA in detecting changes in stem cell aggregates associated with culturing protocols, we performed a comparative analysis between a standard image classifying neural network (ResNet) and a simple feedforward neural network (TDANet) which used our topologically-derived feature set. Frames from a previously published study [10] of time-lapsed microscopy of iPSC aggregates undergoing differentiation from 5 experimental conditions over 48 hours were analyzed for 0th order and 1st order homology; this information was used to train TDANet as described in Materials & Methods. Alternatively, the images were directly fed into ResNet. A model was thus trained, for some particular timepoint, on each image of the 78 colonies (in the case of ResNet) at that timepoint or on each homological data summary of the 77 colonies (in the case of TDANet) from that timepoint (one colony was missing its homological data). See the Processing Coordinate Data section of Materials & Methods for a description of the homological data summary.

We found that both neural networks were successful in classifying the colonies. As expected, the networks tended to perform more poorly at earlier timepoints, when the colonies were mostly pluripotent and thus hadn’t organized their distinguishing features. At later timepoints classification accuracy increased to ∼80% accuracy in the case of TDANet and ∼90% in the case of ResNet. Furthermore, low testing accuracy from a parallel training session where the colonies were randomly assigned a treatment label (see the Training and Randomized Labels section of Materials & Methods) showed the networks were dependent on the underlying biological information. As (C) and (D) of Figure 1 show, on the randomized labels neither TDANet and ResNet could achieve an accuracy little better than randomly choosing one out of five. We conclude that the networks cannot identify sets of colonies whose members are not of the same biological type. Thus, the accuracy shown when trained on the colonies’ correct labels shows each model is extracting information related to the biology of the cell, and not incidental patterns that appear across multiple treatment types.

**Fig 1.**
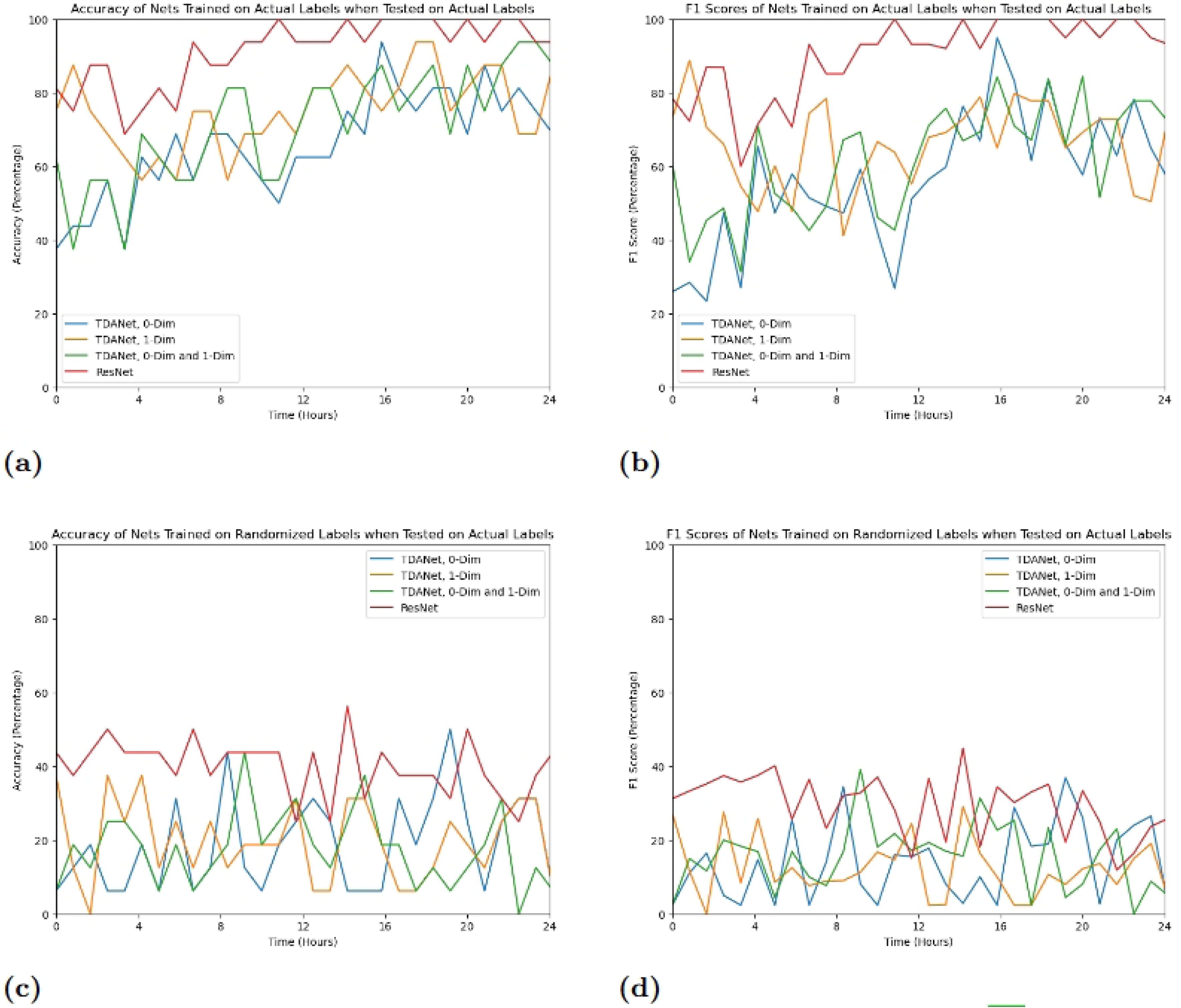
Validation Results of TDANet and ResNet on hiPSC data from [10]. For each timepoint, a neural network model was trained on data from the hiPSCs at that timepoint. The trained model was then tested on a separate set of data from that timepoint that was not used in training. The above graphs show the accuracy of predictions on this validation dataset (A and C), as well as the averaged F1 score (B and D). Results are shown for both the case when a model was trained and tested on both the correct labelings of the colonies (A and B) and on a randomized re-labeling of the colonies (C and D). The three variations of TDANet all depended on the type of training data used: 0-dimensional homology data, 1-dimensional homology data, or a combination of both.

While ResNet tends to have better results than TDANet, both networks exhibit the same behavior. In particular, both networks’ accuracies are characterized by two plateaus. Between timepoints *t*001 and *t*100 (approximately 8.3 hours after beginning to record), the accuracy hovers around a score of ∼60% for TDANet and ∼80% for ResNet, before jumping to their respective optimized rates during the interval of *t*100 to *t*150 (approximately 12.5 hours into imaging). This interval is of interest since it suggests that both networks are sensitive to cell differentiation in this experimental window. That is, the timepoints during which accuracy increases from the lower plateau to the higher plateau might be the window during which the colonies differentiate from their pluripotent and less distinguishable beginnings into their final tissue fates. Past work has shown that information about cell fates appears well before differentiation occurs. For example, reading gene expression levels during pluripotency could predict the percentage of cells that differentiated into cardiomyocytes [11] and hepatocytes [12]. Our work suggests that these clues can be seen in the morphology as well.

An unexpected result of TDANet was the impact of the dimension of the homology data. We had expected TDANet to perform best when given an input of both 1-dimensional and 0-dimensional homology. However, the accuracy of TDANet when trained on this comprehensive data does not significantly exceed its accuracy when given homology data from just one of the dimensions (see Figure 1). In particular, TDANet trained on both 0-dimensional and 1-dimensional homology data does not outperform TDANet trained on just the 1-dimensional data. While the model using both dimensions only takes the first twenty persistence landscape functions for each dimension instead of the first forty, this does not appear to negatively impact its performance. Indeed, we created a version of TDANet that would just take the first twenty persistence landscape functions of a single homology dimension (see Figure 2). We found that the performance of this model approximated the version that used forty persistence landscape functions (Figure 1), suggesting that the critical knowledge for classification resides in the initial landscape functions. As an alternate explanation, we believe that much of the same information about the colony is encoded in both dimensions, so including the persistence landscape functions of more than one is redundant.

**Fig 2.**
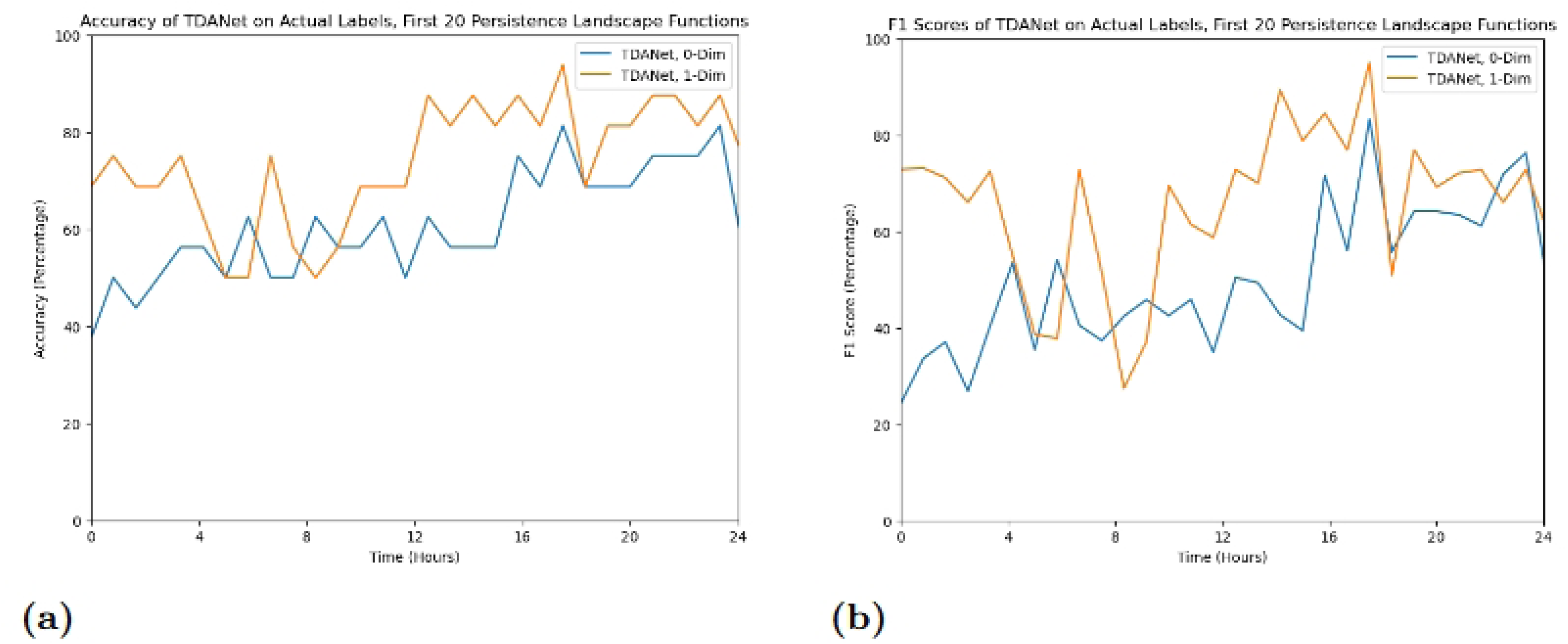
Validation accuracy (A) and F1 scores (B) of TDANet models trained using just the first twenty persistence landscape functions of a single homology dimension.

### Attention analysis suggests topological data can improve use of biological features for ResNet classification

Class activation maps (CAMs) are a useful tool that allow further insight into the decisions made by an image classifier like ResNet (see [13] for a comprehensive explanation). The CAM acts as a heatmap laid over the original image, showing the attention paid by the network to the different regions in the image. Regions that have high values are those that make a larger contribution to the net’s classification decision. In evaluating the CAMs, one can thus determine what parts of the colony are the most impactful in distinguishing it from colonies of different treatment types. Furthermore, the CAM can also allude to whether the network is making the categorization using the “correct” data. That is, if high attention is paid to regions with cells, then this suggests the network is making its decisions based on the underlying biological phenomena in the image. On the other hand, if the heatmap tends to focus on areas that have no cells, this implies the network is using details specific to the image, like the location of the colony in the photo.

In general, the CAMs reveal that ResNet generally focuses on the region of the image that contains the cells. This can be ascertained via a side-by-side comparison of ResNet models with different training protocols, some examples of which can be seen in Figure 3. More CAMs, including movies showing the evolution of CAMs over time for a colony, can be found at https://github.com/aruysdeperez/TDANet.git. The movies show that in contrast to the ResNet models that were trained on randomized labels, the models trained on the correct labels are much more consistent with respect to their distribution of attention.

**Fig 3.**
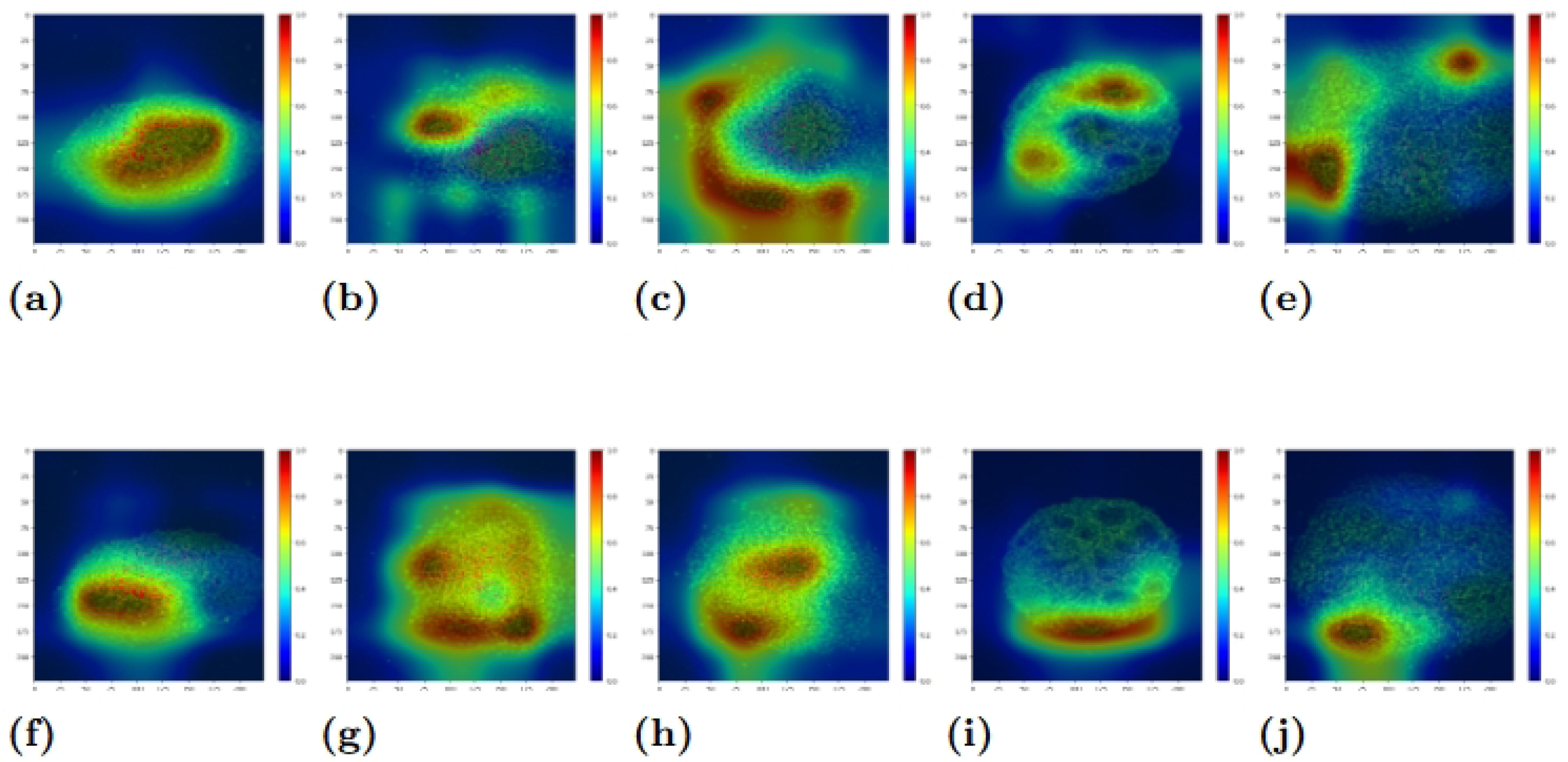
Examples of class activation maps of the five different treatment protocols. Top row (A,B,C,D,E) is the image created by the network trained on the actual labels, while the bottom row (F,G,H,I,J) consists of the same images trained on randomized labels. The columns are arranged by protocol treatment. In order they are WT (A and F), BMP4 (B and G), CHIR (C and H), DS (D and I), and DS+CHIR (E and J).

Each of the different protocol treatments has its own patterning in the class activation map. In the case of the WT and DS-treated colonies (see the Data section of Materials & Methods for a description of the different treatments), the high-value region of the class activation map is centered on the interior of the colony. The difference between these two types lies in the spread of attention across the colony. For the WT colonies, almost the entire aggregate of cells has an activation value of 0.4 or more. However, in the case of the DS colonies, there are large areas of the interior that have an activation value close to 0.

In the case of the BMP4 and CHIR colonies, the CAMs tend to place their attention on the border region of the colony, and place less significance on the interior. This pattern can still be evident of the net’s receptivity to the colony structure: Both the BMP4 and CHIR treated colonies are characterized by a “fringe”, where the cells on the boundary are less densely distributed than the cells in the interior. We conclude that the CAM’s attention to the border demonstrates awareness and use of this feature in the classification process.

For the DS+CHIR colonies, attention tends to focus on the corners of the image. In other colonies these corner locations would have little to no cells, but due to the size of the DS+CHIR colonies they are populated. Thus we concluce that the network is characterizing the DS+CHIR colonies using its larger size, as only this treatment type would show a dense cell population in these areas.

A particularly striking conclusion about the CAMs for the DS treated colonies is that the selected areas the network does pay significant attention to are not necessarily the areas characterized by the “rosettes”. These low density holes that appear in the colony are eye-catching to a human observer and are a distinguishing feature for this type of treatment, as no other colony type shows this motif. It is a surprise then, that the network does not give much importance to these formations. However, in this shortfall we can see an opportunity for topological data analysis. The rosettes are a feature that would stand out in a barcode. The disregard paid them by ResNet shows that there is some relevant information in the topology that is not used by an image classifier.

In summary, the class activation maps reveal that ResNet uses information about the colonies for its classification of the images, and is able to distinguish colonies based on unique biological features. However, as shown in the maps of the DS+CHIR colonies, ResNet can obfuscate the biological data with circumstantial details. Also, as evidenced by the DS colonies, not all relevant details are fully picked up by ResNet. Thus, it appears as though ResNet could benefit from a more directed focus using topological information.

### Networks show similar performance for time-separated training and testing datasets

Another method to evaluate the networks’ robustness involves using datasets from separate times for training and testing. For the results shown in Figure 1 we took training and testing data from the same timepoint. Here, we instead train a model on colony data from *T*, then for a different time *S* ≠ *T*, we have the *T* -trained model classify the data from *S*.

The point of this exercise is to see how well insights made by the networks about the colonies extrapolate over time. We expect that the model trained on data from timepoint *T* will perform better on data from timepoints that are close to *T* compared to timepoints that are further away. We would like to see how quickly this performance decays; that is, how far away a timepoint from *T* can a model trained on *T* sustain high accuracy.

As Figure 4 shows, we do see a trend of models performing better on data from timepoints close to the timepoint on whose data they were trained. This nearby performance effect tends to remain stronger for models trained at later timepoints.

**Fig 4.**
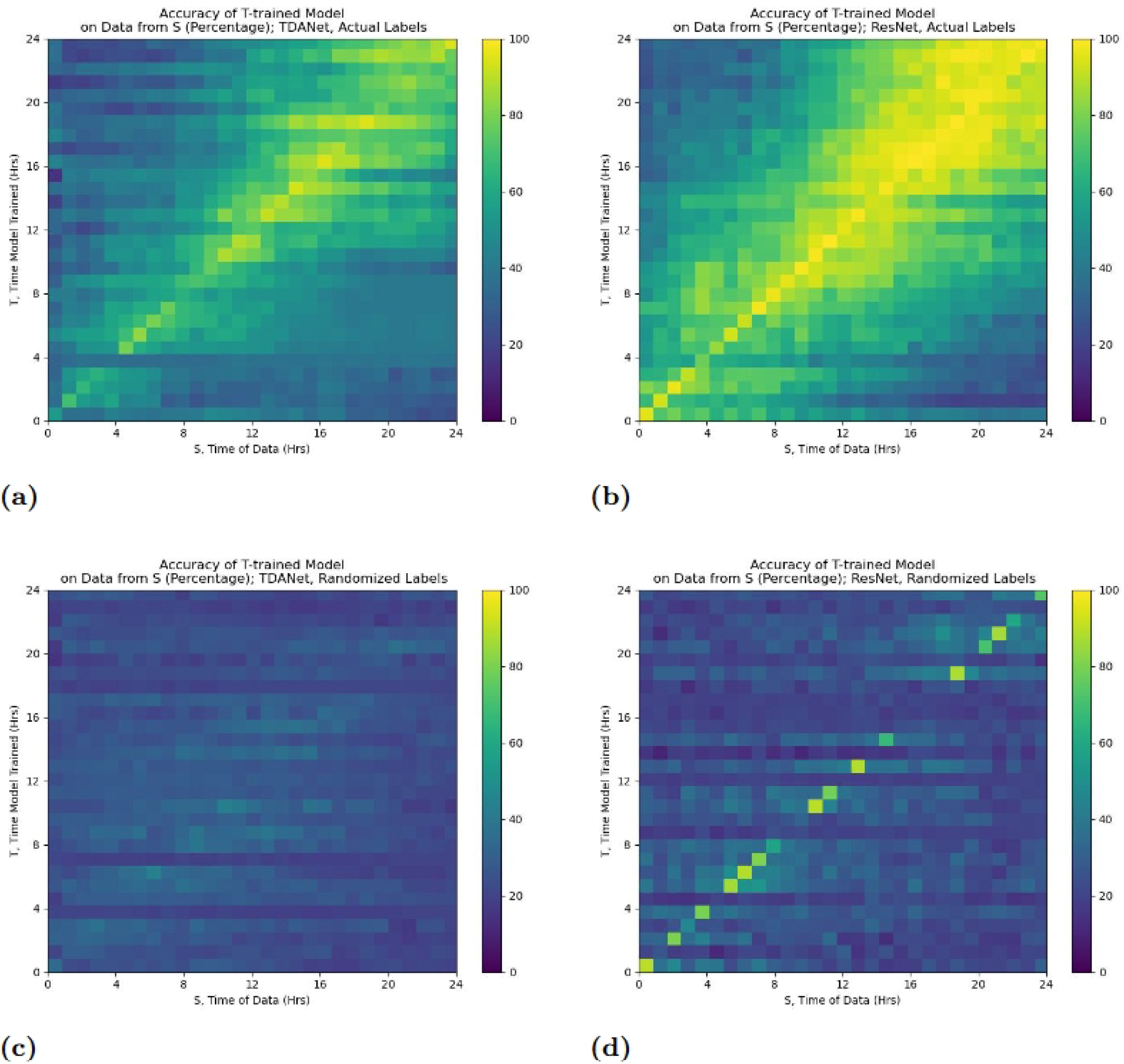
Accuracy of a neural network trained at timepoint *T* and tested on data from a different timepoint *S*. Training was done using both the actual labels (A and B) and the randomized labels (C and D) although testing was always done on the actual labels. Results are for both TDANet (A and C) and ResNet (B and D).

However, an issue with Figure 4 is that it masks the decay with its overall performance. It is possible that some models maintain robust performance with little decay in accuracy that is unseen because they have a lower starting accuracy. To address this we introduce an accuracy metric which we call the *time differential accuracy metric* ⟨*T, S*⟩ for two timepoints *T* and *S*. We define it as

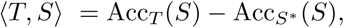

where Acc*_A_*(*B*) is the accuracy of a network trained on data from time *A* when tested on data from time *B*, and *S*^∗^ is a time close to *S* (in our case 50 minutes earlier). The goal of this metric is to show the relative change in accuracy of the neural network model, as this better represents how much accuracy a network can preserve. We choose *S*^∗^ instead of *S* since using an *S*-trained model would suffer from one of two issues: either the testing dataset would include training data it had already seen, or with colony data split into disjoint training and testing sets it would thus be tested on a smaller dataset than the *T* -trained model. The results of the analysis using this metric are in Figure 5.

**Fig 5.**
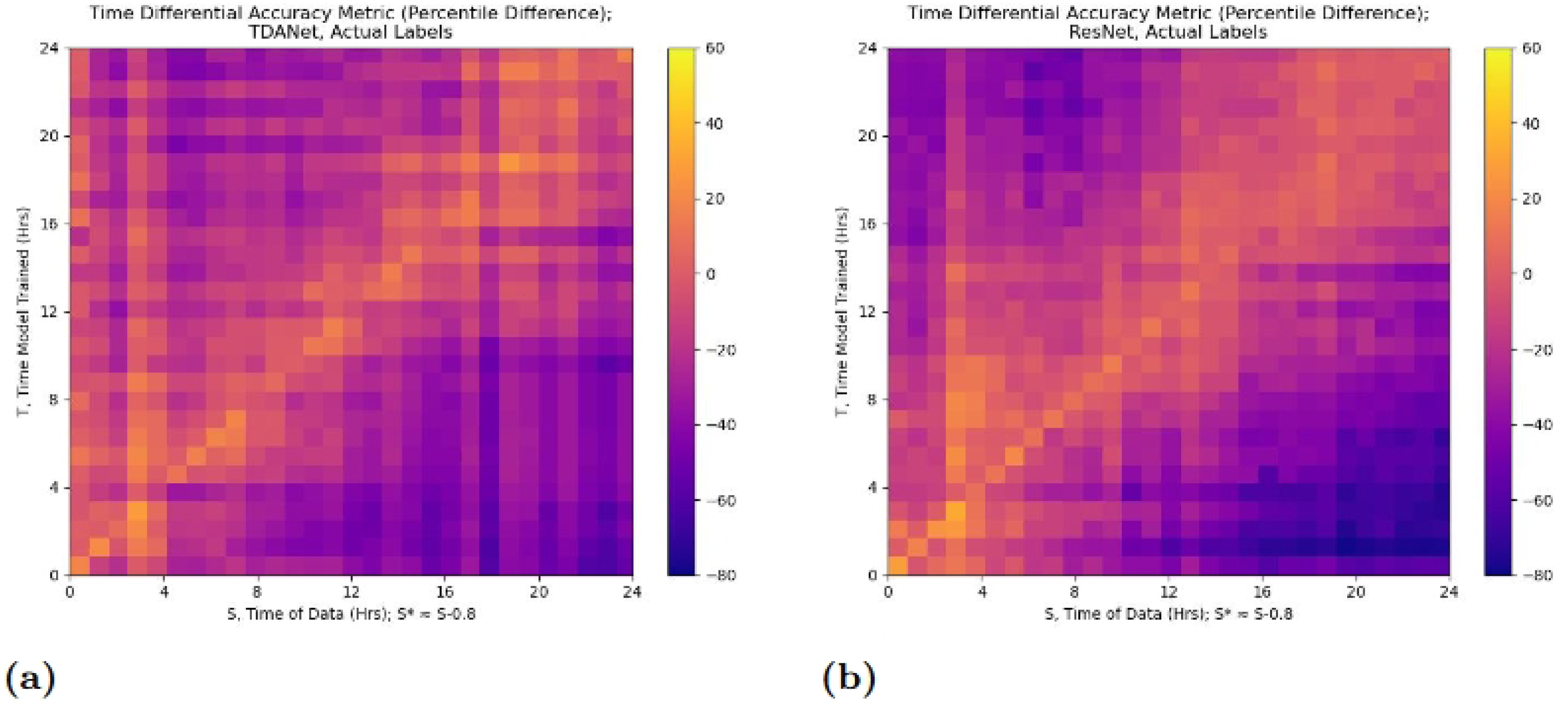
Time differential accuracy metric ⟨*T, S*⟩ for distinct timepoints *S* and *T*. A neural network is trained on data from *T*, along with another network trained on data from *S*^∗^, a time close to *S*. The resulting metric is the difference in accuracy between the *T* -trained network and the *S*^∗^-trained network on data from *S*. Results are for both TDANet (A) and ResNet (B) trained on the actual labels.

There are two observations to be made from the analysis concerning the rate at which model accuracy decays the further the testing time is from the training time. First, for both ResNet and TDANet, the decay rate is slower when the model is trained on data from later timepoints (the high values for the metric at early values of *S* are likely due to the poorer starting accuracy of the *S*^∗^-trained model, as evidenced by the fact that values remain high for all *T*). Second, the decay rate is not appreciably different between ResNet and TDANet. The first result is expected for reasons similar to why we would expect higher accuracy at later timepoints: by these points the colonies have differentiated, and so distinguishing features would have emerged, and which can be seen across many timepoints. The second result is more ambiguous, though it seems to suggest that ResNet and TDANet have equal capability in transferring their insights across time. Perhaps this effect is due to limitations in similarity to colony features at different timepoints.

## Discussion

We investigate the potential for neural networks to accurately classify stem cell fates using morphological data. To do so, we take pluripotent stem cell colonies given one of five differentiation protocol treatments, and ask a neural network to guess the correct protocol. We compare two different network models, each with its own type of information used as input. One is ResNet, a traditional image classifier that uses a photo of the colony as input. The other is TDANet, a simple 4 layer feedforward network that uses topological information in the form of persistent homology created from approximations of the cells’ locations.

Results show that TDANet can classify stem cell colonies with a performance far better than random chance. Network performance is in keeping with assumptions about classification difficulty due to colony differentiation, with networks trained on data from later timepoints tending to have higher accuracy rates. Furthermore, accuracy results reveal two time periods during which accuracy is stable, with a transition occurring roughly 8 to 10 hours after starting data collection. This transition, which can also be observed in the ResNet models, could be the window during which the colonies undergo differentiation, with the earlier time period being when the cells were mostly in their undifferentiated state, and the post-transition period when the colony has achieved its final state.

What is significant about the transition period is that it covers timepoints where many of the colonies’ images are still indistinguishable to the human eye. If indeed this transition is the period of differentiation, then this shows computer vision’s ability to detect structural differences in tissues that is not apparent to humans. Unfortunately, we cannot verify the moment of differentiation from colonies of this particular dataset. However, a straightforward follow-up experiment where differentiation status can be tracked and recorded could ascertain this, and provide a crucial insight.

When compared to the results of ResNet, our TDANet clearly had lower accuracy. However, there are a couple of caveats to simply dismissing TDANet in favor of an image classifier. For one, as our analysis on the class activation maps alludes to, each network has its own set of insights used in the decision making process to which the other does not have access. Furthermore, given the difference in sophistication between the two different neural nets used, it would be inaccurate to conclude that the visual information is superior to the topological. It seems more likely that the performance gap is due to network architecture, and not data input. This leaves open a couple of doors to future work. For one, the performance of the topological method could be significantly improved if we used a more complex network. We ask whether TDANet can be modified or refined into a network that makes use of the persistent homology in a superior manner. Alternatively, topological data could be incorporated into an image classifier. In this conception the image classifier would be told to focus on those parts of the image that have significant homological elements, either through restricting the neural network to those regions or giving them greater weight during decision making.

## Materials and Methods

### Persistent homology

The mathematical tool we use in preparing the colony data is persistent homology. Here, we will present a simplified version for analysis of 2-dimensional data. Those wishing to learn more about the generalized approach should see Edelsbrunner et al. [14] Informally, the job of persistent homology is to provide quantitative data for properties of data sets that are usually described qualitatively. As an example, we begin with the point cloud *X* shown in Figure 6a. At a glance, the data takes the shape of what looks like a barbell, with two separate rings of points joined together. But how do we rigorously describe this? How can we make a case that the point at the very bottom of the figure is “close enough” to complete the lower left circle? How do we make the case that smaller rings, like a triangle made up of three of the points, are not as significant?

**Fig 6.**
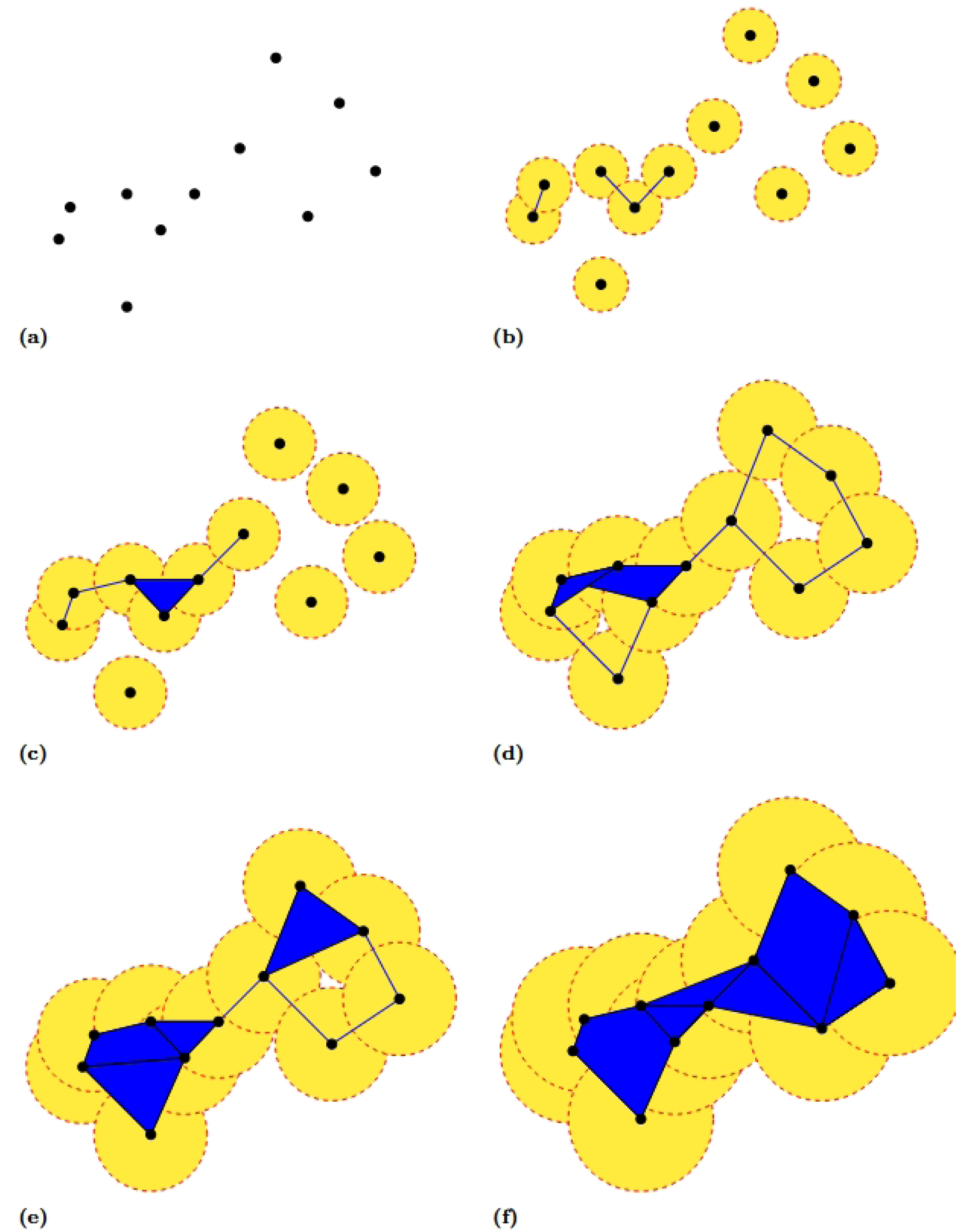
Stages of the growth of the Rips complex for a set of points *X* for increasing values of the radius of the circles used in computing the complex.

We draw a circle of radius *r*, where *r* is a parameter value, around each of the points. We start with a value for *r* so small that none of the circles intersect, and then observe the intersections of the circles as *r* increases.

When *r* becomes large enough such that two circles intersect, we place a line segment between the points for which those circles are the center. In this way we are keeping track of the total number of connected components. In this example, we begin with 11 different components, the points themselves. As points are joined by line segments, the number of components shrinks as they merge together. In Figure 6b, for example, the radius *r* of the circles is large enough that we place down three line segments.

In addition to placing line segments as *r* increases, we also place down polygons. Whenever we have a set of three or more points such that the pairwise intersection of any two of their circles is nonempty, we set down a polygon whose vertices are precisely those points. We can observe an example of this in Figure 6c. For the three points that were joined together in Figure 6b, their circles have now grown large enough that the intersection of any two is nonempty. We thus “fill in” the convex area between the three points to create a triangle. We continue to keep track of the components, noting that at this stage there are 6 components. With the addition of polygons, we are building what is called a *simplicial complex*. We call the simplicial complex constructed in this manner a *Rips complex*.

As we monitor the number of components, we also take note of instances where “holes” appear in our complex. One can see two examples of this in Figure 6d. By this point, the radii have grown large enough that all the points have been connected into one large component. However, the radii are not large enough that we have filled in all the convex areas between points with polygons. As such, the Rips complex has formed two loops that encircle regions of the plane not yet added to the complex. We keep track of the *persistence* of each of these holes through two values of *r*: the value at which it forms and the value at which it is filled in. As one can observe in Figure 6e, there is a value of *r* at which the bottom left has now been filled in, while the top right hole persists, albeit at a smaller size. It is not until a larger value of *r*, shown in Figure 6f do we have that this hole vanishes. At this point our Rips complex is a single connected component without any holes, and will remain as such as *r* continues on to infinity.

We summarize our analysis in a *barcode*, a collection of intervals with each interval representing the lifespan of a particular homology element (a connected component or a hole). The connected components are the *0-dimensional* elements and the holes are the *1-dimensional* elements. The interval’s endpoints are the “birth” and “death” of the corresponding element. These are respectively the value of *r* at which the element comes into being (*r* = 0 for the components; the *r* that connects the loop for the holes) and the value of *r* at which the element ceases to exist (the *r* at which the component merged with another or the hole is completely filled in). The barcode can be represented graphically as a *persistence diagram* [14], as shown in Figure 7.

**Fig 7.**
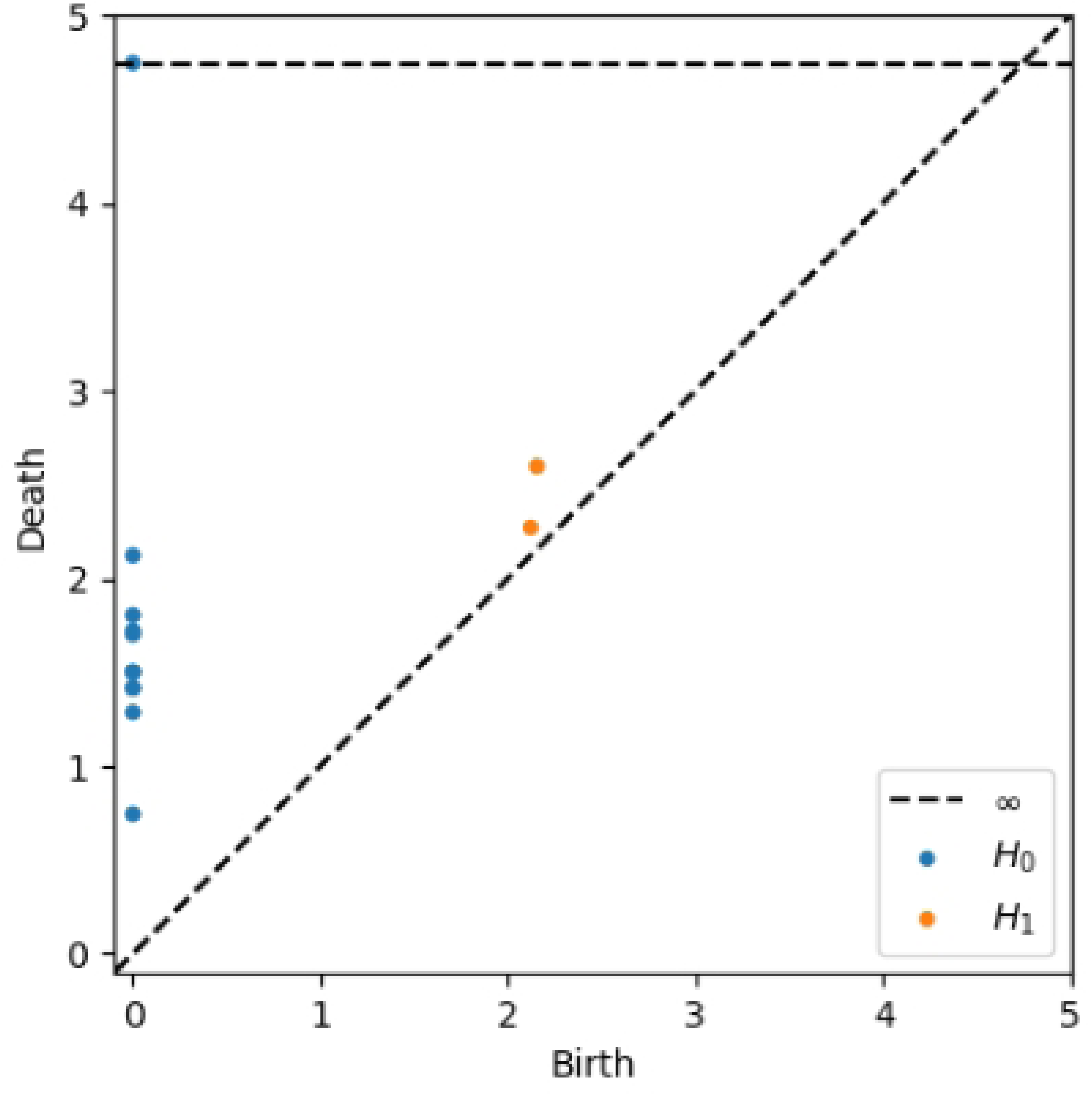
The persistence diagram for the point cloud *X* shown in Figure 6, computed using software from [15]. The blue points represent the connected components (0-dimensional homology elements, or *H*_0_) of the Rips complex. The orange points represent the holes (1-dimensional homology elements, or *H*_1_). The x-axis, labeled “Birth”, denotes the value of *r* at which the element appears (note this is why all the connected components are at *x* = 0, as they existed in the form of individual points at the very beginning). The y-axis, labeled “Death”, denotes the value at which the element disappears. In the case of the components, this is the value at which it connects to another component. For the holes, this is the value at which the hole is “filled in”. The blue point on the dotted line labeled ∞ is the one large component into which all the components are eventually joined.

**Fig 8.**
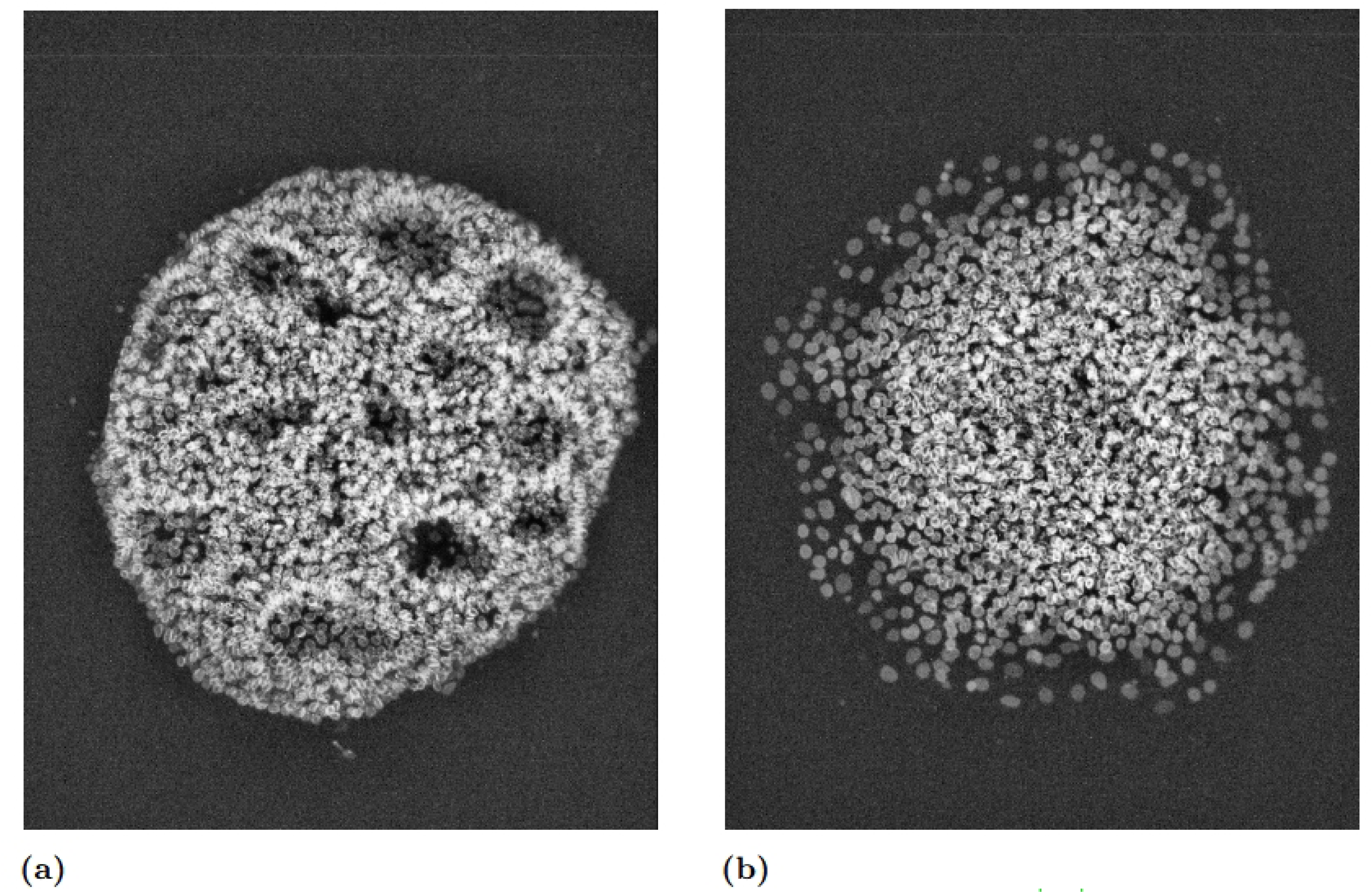
Examples of microscopic images of stem cell colonies from [10], showing rosettes when treated with Dual SMAD (A) and a fringe when treated with BMP4 (B), recorded at timepoint ‘t288’ (approximately 24 hours after starting recording). We found that in many of the colonies, distinctive structural features appear at later times. Two in particular stand out: the rosettes in colonies treated with dual SMAD, and a fringe around the outer edges of the colony that appears in BMP4 and CHIR treated colonies.

With the barcode, we now have concrete metrics about the density and clustering of the data set and can thus elucidate its shape more clearly. Returning to the beginning of our example, we can see that we are justified in our characterization of *X* as two rings joined together. The barcode has precisely two 1-dimensional homology elements; the two rings we originally saw at first glance.

The approach we have detailed in the example is the one we take with our colony data. Like the data set *X* of the example, our raw data consists of points in the coordinate plane. Each data set consists of a single colony at a particular timepoint, and each point represents an approximation of the location of a cell nucleus. We construct a Rips complex and record the values of *r* at which the components and holes of the complex appear and disappear during its construction. We use the program Ripser for the computation of the complexes and the resulting barcodes [15].

### Data

We use data on 78 human induced pluripotent stem cell (hiPSC) colonies from the work of [10] as provided by the authors. Each of these colonies was either left as wild type (WT) or treated with one of four morphogen combinations: BMP4, dual SMAD inhibition (DS), CHIR, or combined dual SMAD inhibition and CHIR (DS+CHIR).

Each of these morphogens affects differentiation during gastrulation. CHIR is an activator of the WNT pathway [16]. Suppression of BMP4 has been shown to lead to the failure of the mesoderm to develop [17], and SMAD features in the Nodal pathway, which promotes mesoderm formation [18] [19].

Every colony was imaged over a 24-hour period. Images of each colony were acquired every 5 minutes, for a total of 288 individual frames. The images, which were 834 × 1094 pixel photos of a 76.50 × 1002.83 micron space, were then analyzed for cell detection and segmentation by the authors to create a list of coordinates approximating the locations of each of the cells, which tended to number between 500 and 2000 per colony. Thus, each colony has 288 sets of coordinate data derived from 288 images, yielding 78 time series. For each protocol treatment, the colonies were created by first aggregating the cells over a 24-hour period, after which the aggregates were plated down in culture wells and developed into colonies. In the case of the BMP4 and CHIR treatments, there was a 24-hour wait period, after which the ligands were applied for another 24 hours before the final 24-hour imaging process. In the case of the DS treatment, the treating of the cells occurred at the beginning of the aggregation period, and continued throughout the consecutive 24-hour periods of aggregation, post-seeding colony formation, and imaging. For the DS+CHIR treatment, CHIR was applied 48 hours before aggregation, and was maintained through the 48 hour pre-aggregation period, the 24 hour aggregation period, the 24 post-seeding period, and the 24 hour imaging period [10].

### Processing Coordinate Data

We used Ripser [15] to create a barcode from the coordinate data of a colony at a particular timepoint.

However, using a barcode as input to our feedforward neural network raises the difficulty of fitting a data structure of varying length to a fixed input size parameter. We wanted to avoid imposing an arbitrary cutoff on the number of intervals from a barcode given to a network. Thus, we implemented a new input format using the *persistence landscape*.

We define the *persistence landscape* [20] as follows. For an interval (*b, d*) in the barcode, define the function *f*_(_*_b,d_*_)_ : [0, ∞) → [0, ∞) to be

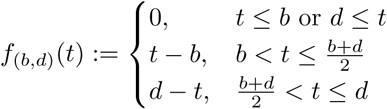

The persistence landscape is then a family of functions 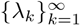 where the *k*th persistence landscape function is given by

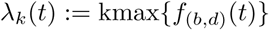

over all persistence elements (*b, d*) in the barcode. Here, *kmax* means the *k*th largest element in the set.

Informally, the persistence landscapes are used here to find the long-lasting features, but in the context of comparing with features that occur for the same values of *r*. For example, say a barcode has an interval [*r*_0_*, r*_1_). Suppose that this interval, while shorter than many other elements of the barcode, is the longest such interval among intervals that extend into the region between *r*_0_ and *r*_1_. Then this interval would feature prominently in the persistence landscapes, as while it is not a large interval, it is a large interval in its locality.

We selected as sample points for the filtration parameter *r* the values 1, 2*, . . .,* 40 and the first forty persistence landscape functions *λ*_1_*, λ*_2_*, . . ., λ*_40_. We decided on the number 40 in both cases based on examination of the persistence landscape functions of the colonies at various timepoints, which found little information for parameter values greater than 40 or for *λ_i_*when *i >* 40 (see Figure 9 as an example). The input to the neural network is thus a 40 × 40 matrix whose (*i, j*)th entry is *λ_i_*(*j*).

**Fig 9.**
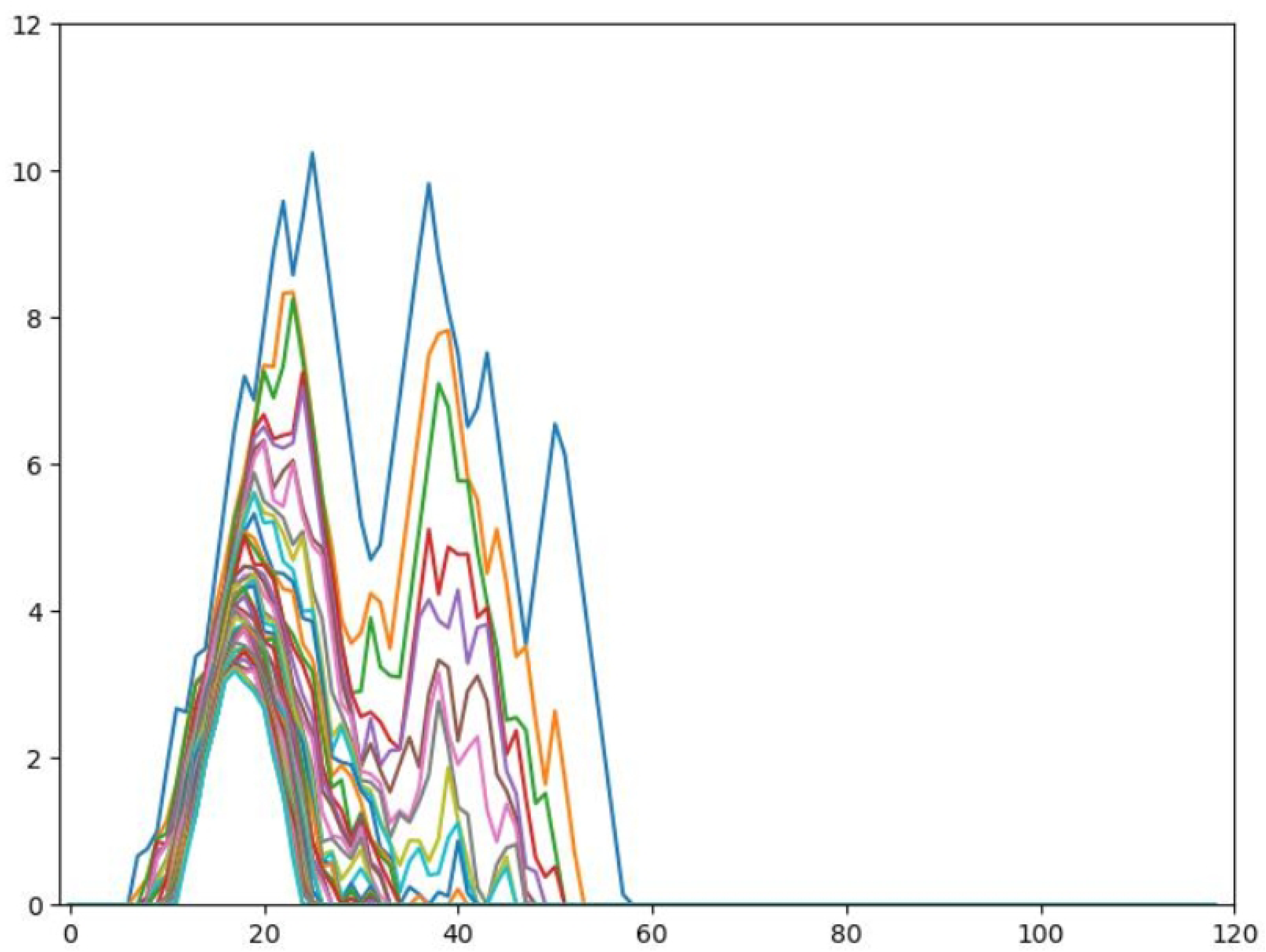
A graph of the first fifty persistence landscape functions of the persistent homology of a stem cell colony.

### TDA-Using Neural Network

We used a feedforward neural network, which we call *TDANet*, to interpret the colonies’ homology data. The network consists of three dense 20-neuron hidden layers with the ReLu activation, and a 5-neuron output layer with the Softmax activation function. The output consists of a “vector of probabilities”, with each index corresponding to a treatment type, and whose entry represents the likelihood the machine has that the input colony is of that type. We used categorical cross-entropy as our loss function, which is given by

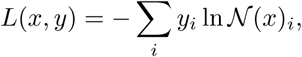

where N() represents the output of the network and (*x, y*) is an input-output pair. We used the ‘adam’ optimizer. Our code for the models, as well as processing the coordinate data into barcodes and then persistence landscapes, can be found on Github: https://github.com/aruysdeperez/TDANet.git. See Figure 10a for a conceptual diagram of TDANet.

**Fig 10.**
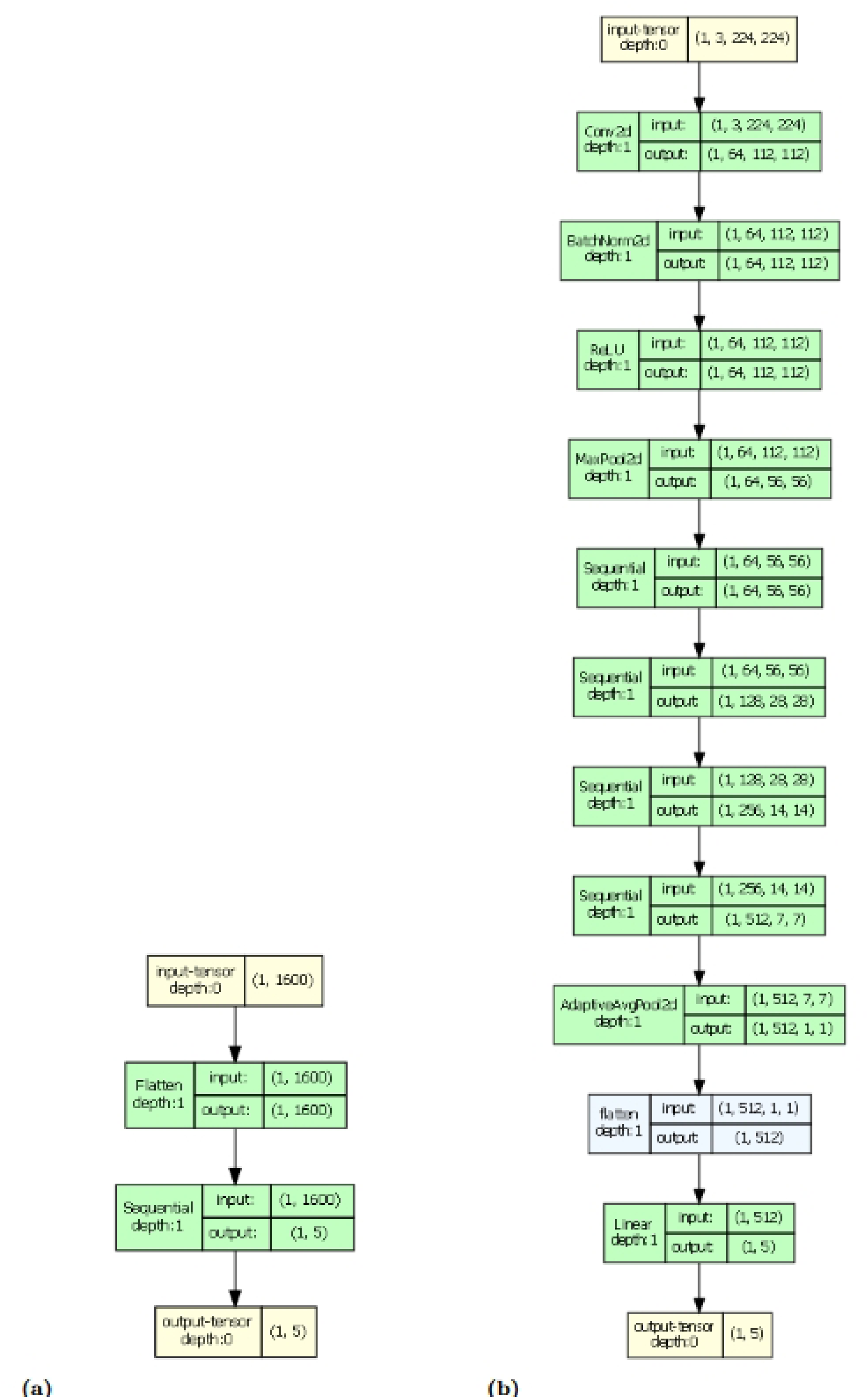
Conceptual diagrams detailing the architecture of TDANet (A) and ResNet (B). For TDANet the input is a 40 × 40 matrix of persistence landscape values for a stem cell colony at a particular timepoint, flattened into a one-dimensional vector. Output is a vector of 5 entries detailing the likelihoods the input colony was a protocol treatment. (B) For ResNet, we changed the standard ResNet architecture so that its output layer was replaced with the same output layer for TDANet. Graph model made using torchview package [22].

### Convolutional Neural Network

To compare our method with more traditional image classifiers, we also trained the convolutional neural network ResNet on the images of the colonies. This model is the 18-layer pretrained version provided by PyTorch [21]. We replaced the output layer with one classifying our five treatment categories. For training this model, we froze all the parameters except for our new output layer, so only that layer would be updated. See Figure 10b for a conceptual diagram of ResNet.

We trained both of our neural networks on a timepoint by timepoint basis. That is, we fixed a timepoint and trained solely on colony data from that particular timepoint. Our justification is that the network’s performance is based on time elapsed since treatment: the network will be more successful at distinguishing colonies at later timepoints, as these will be more differentiated. Thus, in training timepoint by timepoint we could establish a graded measure of the network’s accuracy based on the differentiation of the cells.

### Training and Randomized Labels

For both neural networks, we divided the timepoint data such that 80% would be used for training, and the rest for validation. We trained both networks for 50 epochs.

To measure the significance of the biological classifications, we also performed training on the colonies with randomized labels. That is, instead of giving the correct treatment label to a colony, we used a label that was randomly chosen from one of the five possible. See Table 1 for the randomized distribution of labels. The training of the randomized labels also underwent an 80-20 split, although during testing we used the actual labels on the testing data set rather than the randomized ones.

**Table 1.** Confusion matrix for the randomized label set used to control for the performance of TDANet and ResNet. The (*i, j*)th entry represents the number of colonies whose actual label was *i*, but were assigned label *j*. *There was one BMP4 colony randomly labeled as DS+CHIR, for which we had colony images but lacked the cell coordinate data. Thus, while we could use this colony for training and testing ResNet, it was not available for use by TDANet.

|  |  | Label Randomly Assigned |  |  |  |  |
| --- | --- | --- | --- | --- | --- | --- |
|  |  | BMP4 | CHIR | DS | DS+CHIR | WT |
| Actual<br>Label | BMP4 | 2 | 1 | 4 | 4(3)* | 5 |
|  | CHIR | 3 | 3 | 3 | 3 | 3 |
|  | DS | 3 | 3 | 1 | 3 | 4 |
|  | DS+CHIR | 3 | 2 | 4 | 3 | 5 |
|  | WT | 2 | 2 | 2 | 3 | 3 |

## Acknowledgments

We thank David Joy and the Todd McDevitt lab for their sharing of the colony images and data.

## Supporting information

**S1 CAMActual 22 01 WT.** CAM Activation Maps of Colony 22 01 (WT Treatment) Using Model Trained on Actual Labels

**S2 CAMActual 22 02 WT.** CAM Activation Maps of Colony 22 02 (WT Treatment) Using Model Trained on Actual Labels

**S3 CAMActual 22 03 WT.** CAM Activation Maps of Colony 22 03 (WT Treatment) Using Model Trained on Actual Labels

**S4 CAMActual 22 04 WT.** CAM Activation Maps of Colony 22 04 (WT Treatment) Using Model Trained on Actual Labels

**S5 CAMActual 22 05 WT.** CAM Activation Maps of Colony 22 05 (WT Treatment) Using Model Trained on Actual Labels

**S6 CAMActual 22 06 WT.** CAM Activation Maps of Colony 22 06 (WT Treatment) Using Model Trained on Actual Labels

**S7 CAMActual 22 07 WT.** CAM Activation Maps of Colony 22 07 (WT Treatment) Using Model Trained on Actual Labels

**S8 CAMActual 22 08 WT.** CAM Activation Maps of Colony 22 08 (WT Treatment) Using Model Trained on Actual Labels

**S9 CAMActual 22 09 WT.** CAM Activation Maps of Colony 22 09 (WT Treatment) Using Model Trained on Actual Labels

**S10 CAMActual 22 10 WT.** CAM Activation Maps of Colony 22 10 (WT Treatment) Using Model Trained on Actual Labels

**S11 CAMActual 22 11 WT.** CAM Activation Maps of Colony 22 11 (WT Treatment) Using Model Trained on Actual Labels

**S12 CAMActual 22 12 WT.** CAM Activation Maps of Colony 22 12 (WT Treatment) Using Model Trained on Actual Labels

**S13 CAMActual 22 13 BMP4.** CAM Activation Maps of Colony 22 13 (BMP4 Treatment) Using Model Trained on Actual Labels

**S14 CAMActual 22 14 BMP4.** CAM Activation Maps of Colony 22 14 (BMP4 Treatment) Using Model Trained on Actual Labels

**S15 CAMActual 22 15 BMP4.** CAM Activation Maps of Colony 22 15 (BMP4 Treatment) Using Model Trained on Actual Labels

**S16 CAMActual 22 16 BMP4.** CAM Activation Maps of Colony 22 16 (BMP4 Treatment) Using Model Trained on Actual Labels

**S17 CAMActual 22 17 BMP4.** CAM Activation Maps of Colony 22 17 (BMP4 Treatment) Using Model Trained on Actual Labels

**S18 CAMActual 22 18 BMP4.** CAM Activation Maps of Colony 22 18 (BMP4 Treatment) Using Model Trained on Actual Labels

**S19 CAMActual 22 19 BMP4.** CAM Activation Maps of Colony 22 19 (BMP4 Treatment) Using Model Trained on Actual Labels

**S20 CAMActual 22 20 BMP4.** CAM Activation Maps of Colony 22 20 (BMP4 Treatment) Using Model Trained on Actual Labels

**S21 CAMActual 22 21 BMP4.** CAM Activation Maps of Colony 22 21 (BMP4 Treatment) Using Model Trained on Actual Labels

**S22 CAMActual 22 23 BMP4.** CAM Activation Maps of Colony 22 23 (BMP4 Treatment) Using Model Trained on Actual Labels

**S23 CAMActual 22 24 BMP4.** CAM Activation Maps of Colony 22 24 (BMP4 Treatment) Using Model Trained on Actual Labels

**S24 CAMActual 22 25 BMP4.** CAM Activation Maps of Colony 22 25 (BMP4 Treatment) Using Model Trained on Actual Labels

**S25 CAMActual 22 26 BMP4.** CAM Activation Maps of Colony 22 26 (BMP4 Treatment) Using Model Trained on Actual Labels

**S26 CAMActual 22 27 BMP4.** CAM Activation Maps of Colony 22 27 (BMP4 Treatment) Using Model Trained on Actual Labels

**S27 CAMActual 22 28 BMP4.** CAM Activation Maps of Colony 22 28 (BMP4 Treatment) Using Model Trained on Actual Labels

**S28 CAMActual 22 29 CHIR.** CAM Activation Maps of Colony 22 29 (CHIR Treatment) Using Model Trained on Actual Labels

**S29 CAMActual 22 30 CHIR.** CAM Activation Maps of Colony 22 30 (CHIR Treatment) Using Model Trained on Actual Labels

**S30 CAMActual 22 31 CHIR.** CAM Activation Maps of Colony 22 31 (CHIR Treatment) Using Model Trained on Actual Labels

**S31 CAMActual 22 32 CHIR.** CAM Activation Maps of Colony 22 32 (CHIR Treatment) Using Model Trained on Actual Labels

**S32 CAMActual 22 33 CHIR.** CAM Activation Maps of Colony 22 33 (CHIR Treatment) Using Model Trained on Actual Labels

**S33 CAMActual 22 34 CHIR.** CAM Activation Maps of Colony 22 34 (CHIR Treatment) Using Model Trained on Actual Labels

**S34 CAMActual 22 35 CHIR.** CAM Activation Maps of Colony 22 35 (CHIR Treatment) Using Model Trained on Actual Labels

**S35 CAMActual 22 36 CHIR.** CAM Activation Maps of Colony 22 36 (CHIR Treatment) Using Model Trained on Actual Labels

**S36 CAMActual 22 37 CHIR.** CAM Activation Maps of Colony 22 37 (CHIR Treatment) Using Model Trained on Actual Labels

**S37 CAMActual 22 39 CHIR.** CAM Activation Maps of Colony 22 39 (CHIR Treatment) Using Model Trained on Actual Labels

**S38 CAMActual 22 40 CHIR.** CAM Activation Maps of Colony 22 40 (CHIR Treatment) Using Model Trained on Actual Labels

**S39 CAMActual 22 41 CHIR.** CAM Activation Maps of Colony 22 41 (CHIR Treatment) Using Model Trained on Actual Labels

**S40 CAMActual 22 42 CHIR.** CAM Activation Maps of Colony 22 42 (CHIR Treatment) Using Model Trained on Actual Labels

**S41 CAMActual 22 43 CHIR.** CAM Activation Maps of Colony 22 43 (CHIR Treatment) Using Model Trained on Actual Labels

**S42 CAMActual 22 44 CHIR.** CAM Activation Maps of Colony 22 44 (CHIR Treatment) Using Model Trained on Actual Labels

**S43 CAMActual 26 01 DS.** CAM Activation Maps of Colony 26 01 (DS Treatment) Using Model Trained on Actual Labels

**S44 CAMActual 26 02 DS+CHIR.** CAM Activation Maps of Colony 26 02 (DS+CHIR Treatment) Using Model Trained on Actual Labels

**S45 CAMActual 26 03 DS.** CAM Activation Maps of Colony 26 03 (DS Treatment) Using Model Trained on Actual Labels

**S46 CAMActual 26 04 DS.** CAM Activation Maps of Colony 26 04 (DS Treatment) Using Model Trained on Actual Labels

**S47 CAMActual 26 05 DS+CHIR.** CAM Activation Maps of Colony 26 05 (DS+CHIR Treatment) Using Model Trained on Actual Labels

**S48 CAMActual 26 06 DS.** CAM Activation Maps of Colony 26 06 (DS Treatment) Using Model Trained on Actual Labels

**S49 CAMActual 26 07 DS+CHIR.** CAM Activation Maps of Colony 26 07 (DS+CHIR Treatment) Using Model Trained on Actual Labels

**S50 CAMActual 26 08 DS.** CAM Activation Maps of Colony 26 08 (DS Treatment) Using Model Trained on Actual Labels

**S51 CAMActual 26 09 DS+CHIR.** CAM Activation Maps of Colony 26 09 (DS+CHIR Treatment) Using Model Trained on Actual Labels

**S52 CAMActual 26 10 DS.** CAM Activation Maps of Colony 26 10 (DS Treatment) Using Model Trained on Actual Labels

**S53 CAMActual 26 11 DS.** CAM Activation Maps of Colony 26 11 (DS Treatment) Using Model Trained on Actual Labels

**S54 CAMActual 26 12 DS.** CAM Activation Maps of Colony 26 12 (DS Treatment) Using Model Trained on Actual Labels

**S55 CAMActual 26 13 DS+CHIR.** CAM Activation Maps of Colony 26 13 (DS+CHIR Treatment) Using Model Trained on Actual Labels

**S56 CAMActual 26 14 DS+CHIR.** CAM Activation Maps of Colony 26 14 (DS+CHIR Treatment) Using Model Trained on Actual Labels

**S57 CAMActual 26 15 DS+CHIR.** CAM Activation Maps of Colony 26 15 (DS+CHIR Treatment) Using Model Trained on Actual Labels

**S58 CAMActual 26 16 DS.** CAM Activation Maps of Colony 26 16 (DS Treatment) Using Model Trained on Actual Labels

**S59 CAMActual 26 17 DS+CHIR.** CAM Activation Maps of Colony 26 17 (DS+CHIR Treatment) Using Model Trained on Actual Labels

**S60 CAMActual 26 18 DS.** CAM Activation Maps of Colony 26 18 (DS Treatment) Using Model Trained on Actual Labels

**S61 CAMActual 26 19 DS+CHIR.** CAM Activation Maps of Colony 26 19 (DS+CHIR Treatment) Using Model Trained on Actual Labels

**S62 CAMActual 26 20 DS+CHIR.** CAM Activation Maps of Colony 26 20 (DS+CHIR Treatment) Using Model Trained on Actual Labels

**S63 CAMActual 26 21 DS+CHIR.** CAM Activation Maps of Colony 26 21 (DS+CHIR Treatment) Using Model Trained on Actual Labels

**S64 CAMActual 26 22 DS+CHIR.** CAM Activation Maps of Colony 26 22 (DS+CHIR Treatment) Using Model Trained on Actual Labels

**S65 CAMActual 26 23 DS+CHIR.** CAM Activation Maps of Colony 26 23 (DS+CHIR Treatment) Using Model Trained on Actual Labels

**S66 CAMActual 26 24 DS.** CAM Activation Maps of Colony 26 24 (DS Treatment) Using Model Trained on Actual Labels

**S67 CAMActual 26 25 DS.** CAM Activation Maps of Colony 26 25 (DS Treatment) Using Model Trained on Actual Labels

**S68 CAMActual 26 26 DS.** CAM Activation Maps of Colony 26 26 (DS Treatment) Using Model Trained on Actual Labels

**S69 CAMActual 26 27 DS+CHIR.** CAM Activation Maps of Colony 26 27 (DS+CHIR Treatment) Using Model Trained on Actual Labels

**S70 CAMActual 26 28 DS.** CAM Activation Maps of Colony 26 28 (DS Treatment) Using Model Trained on Actual Labels

**S71 CAMActual 26 29 DS+CHIR.** CAM Activation Maps of Colony 26 29 (DS+CHIR Treatment) Using Model Trained on Actual Labels

**S72 CAMActual 26 30 DS.** CAM Activation Maps of Colony 26 30 (DS Treatment) Using Model Trained on Actual Labels

**S73 CAMActual 26 31 DS+CHIR.** CAM Activation Maps of Colony 26 31 (DS+CHIR Treatment) Using Model Trained on Actual Labels

**S74 CAMActual 26 32 DS.** CAM Activation Maps of Colony 26 32 (DS Treatment) Using Model Trained on Actual Labels

**S75 CAMActual 26 33 DS+CHIR.** CAM Activation Maps of Colony 26 33 (DS+CHIR Treatment) Using Model Trained on Actual Labels

**S76 CAMActual 26 34 DS.** CAM Activation Maps of Colony 26 34 (DS Treatment) Using Model Trained on Actual Labels

**S77 CAMRand 22 01 WT.** CAM Activation Maps of Colony 22 01 (WT Treatment) Using Model Trained on Randomized Labels

**S78 CAMRand 22 02 WT.** CAM Activation Maps of Colony 22 02 (WT Treatment) Using Model Trained on Randomized Labels

**S79 CAMRand 22 03 WT.** CAM Activation Maps of Colony 22 03 (WT Treatment) Using Model Trained on Randomized Labels

**S80 CAMRand 22 04 WT.** CAM Activation Maps of Colony 22 04 (WT Treatment) Using Model Trained on Randomized Labels

**S81 CAMRand 22 05 WT.** CAM Activation Maps of Colony 22 05 (WT Treatment) Using Model Trained on Randomized Labels

**S82 CAMRand 22 06 WT.** CAM Activation Maps of Colony 22 06 (WT Treatment) Using Model Trained on Randomized Labels

**S83 CAMRand 22 07 WT.** CAM Activation Maps of Colony 22 07 (WT Treatment) Using Model Trained on Randomized Labels

**S84 CAMRand 22 08 WT.** CAM Activation Maps of Colony 22 08 (WT Treatment) Using Model Trained on Randomized Labels

**S85 CAMRand 22 09 WT.** CAM Activation Maps of Colony 22 09 (WT Treatment) Using Model Trained on Randomized Labels

**S86 CAMRand 22 10 WT.** CAM Activation Maps of Colony 22 10 (WT Treatment) Using Model Trained on Randomized Labels

**S87 CAMRand 22 11 WT.** CAM Activation Maps of Colony 22 11 (WT Treatment) Using Model Trained on Randomized Labels

**S88 CAMRand 22 12 WT.** CAM Activation Maps of Colony 22 12 (WT Treatment) Using Model Trained on Randomized Labels

**S89 CAMRand 22 13 BMP4.** CAM Activation Maps of Colony 22 13 (BMP4 Treatment) Using Model Trained on Randomized Labels

**S90 CAMRand 22 14 BMP4.** CAM Activation Maps of Colony 22 14 (BMP4 Treatment) Using Model Trained on Randomized Labels

**S91 CAMRand 22 15 BMP4.** CAM Activation Maps of Colony 22 15 (BMP4 Treatment) Using Model Trained on Randomized Labels

**S92 CAMRand 22 16 BMP4.** CAM Activation Maps of Colony 22 16 (BMP4 Treatment) Using Model Trained on Randomized Labels

**S93 CAMRand 22 17 BMP4.** CAM Activation Maps of Colony 22 17 (BMP4 Treatment) Using Model Trained on Randomized Labels

**S94 CAMRand 22 18 BMP4.** CAM Activation Maps of Colony 22 18 (BMP4 Treatment) Using Model Trained on Randomized Labels

**S95 CAMRand 22 19 BMP4.** CAM Activation Maps of Colony 22 19 (BMP4 Treatment) Using Model Trained on Randomized Labels

**S96 CAMRand 22 20 BMP4.** CAM Activation Maps of Colony 22 20 (BMP4 Treatment) Using Model Trained on Randomized Labels

**S97 CAMRand 22 21 BMP4.** CAM Activation Maps of Colony 22 21 (BMP4 Treatment) Using Model Trained on Randomized Labels

**S98 CAMRand 22 23 BMP4.** CAM Activation Maps of Colony 22 23 (BMP4 Treatment) Using Model Trained on Randomized Labels

**S99 CAMRand 22 24 BMP4.** CAM Activation Maps of Colony 22 24 (BMP4 Treatment) Using Model Trained on Randomized Labels

**S100 CAMRand 22 25 BMP4.** CAM Activation Maps of Colony 22 25 (BMP4 Treatment) Using Model Trained on Randomized Labels

**S101 CAMRand 22 26 BMP4.** CAM Activation Maps of Colony 22 26 (BMP4 Treatment) Using Model Trained on Randomized Labels

**S102 CAMRand 22 27 BMP4.** CAM Activation Maps of Colony 22 27 (BMP4 Treatment) Using Model Trained on Randomized Labels

**S103 CAMRand 22 28 BMP4.** CAM Activation Maps of Colony 22 28 (BMP4 Treatment) Using Model Trained on Randomized Labels

**S104 CAMRand 22 29 CHIR.** CAM Activation Maps of Colony 22 29 (CHIR Treatment) Using Model Trained on Randomized Labels

**S105 CAMRand 22 30 CHIR.** CAM Activation Maps of Colony 22 30 (CHIR Treatment) Using Model Trained on Randomized Labels

**S106 CAMRand 22 31 CHIR.** CAM Activation Maps of Colony 22 31 (CHIR Treatment) Using Model Trained on Randomized Labels

**S107 CAMRand 22 32 CHIR.** CAM Activation Maps of Colony 22 32 (CHIR Treatment) Using Model Trained on Randomized Labels

**S108 CAMRand 22 33 CHIR.** CAM Activation Maps of Colony 22 33 (CHIR Treatment) Using Model Trained on Randomized Labels

**S109 CAMRand 22 34 CHIR.** CAM Activation Maps of Colony 22 34 (CHIR Treatment) Using Model Trained on Randomized Labels

**S110 CAMRand 22 35 CHIR.** CAM Activation Maps of Colony 22 35 (CHIR Treatment) Using Model Trained on Randomized Labels

**S111 CAMRand 22 36 CHIR.** CAM Activation Maps of Colony 22 36 (CHIR Treatment) Using Model Trained on Randomized Labels

**S112 CAMRand 22 37 CHIR.** CAM Activation Maps of Colony 22 37 (CHIR Treatment) Using Model Trained on Randomized Labels

**S113 CAMRand 22 39 CHIR.** CAM Activation Maps of Colony 22 39 (CHIR Treatment) Using Model Trained on Randomized Labels

**S114 CAMRand 22 40 CHIR.** CAM Activation Maps of Colony 22 40 (CHIR Treatment) Using Model Trained on Randomized Labels

**S115 CAMRand 22 41 CHIR.** CAM Activation Maps of Colony 22 41 (CHIR Treatment) Using Model Trained on Randomized Labels

**S116 CAMRand 22 42 CHIR.** CAM Activation Maps of Colony 22 42 (CHIR Treatment) Using Model Trained on Randomized Labels

**S117 CAMRand 22 43 CHIR.** CAM Activation Maps of Colony 22 43 (CHIR Treatment) Using Model Trained on Randomized Labels

**S118 CAMRand 22 44 CHIR.** CAM Activation Maps of Colony 22 44 (CHIR Treatment) Using Model Trained on Randomized Labels

**S119 CAMRand 26 01 DS.** CAM Activation Maps of Colony 26 01 (DS Treatment) Using Model Trained on Randomized Labels

**S120 CAMRand 26 02 DS+CHIR.** CAM Activation Maps of Colony 26 02 (DS+CHIR Treatment) Using Model Trained on Randomized Labels

**S121 CAMRand 26 03 DS.** CAM Activation Maps of Colony 26 03 (DS Treatment) Using Model Trained on Randomized Labels

**S122 CAMRand 26 04 DS.** CAM Activation Maps of Colony 26 04 (DS Treatment) Using Model Trained on Randomized Labels

**S123 CAMRand 26 05 DS+CHIR.** CAM Activation Maps of Colony 26 05 (DS+CHIR Treatment) Using Model Trained on Randomized Labels

**S124 CAMRand 26 06 DS.** CAM Activation Maps of Colony 26 06 (DS Treatment) Using Model Trained on Randomized Labels

**S125 CAMRand 26 07 DS+CHIR.** CAM Activation Maps of Colony 26 07 (DS+CHIR Treatment) Using Model Trained on Randomized Labels

**S126 CAMRand 26 08 DS.** CAM Activation Maps of Colony 26 08 (DS Treatment) Using Model Trained on Randomized Labels

**S127 CAMRand 26 09 DS+CHIR.** CAM Activation Maps of Colony 26 09 (DS+CHIR Treatment) Using Model Trained on Randomized Labels

**S128 CAMRand 26 10 DS.** CAM Activation Maps of Colony 26 10 (DS Treatment) Using Model Trained on Randomized Labels

**S129 CAMRand 26 11 DS.** CAM Activation Maps of Colony 26 11 (DS Treatment) Using Model Trained on Randomized Labels

**S130 CAMRand 26 12 DS.** CAM Activation Maps of Colony 26 12 (DS Treatment) Using Model Trained on Randomized Labels

**S131 CAMRand 26 13 DS+CHIR.** CAM Activation Maps of Colony 26 13 (DS+CHIR Treatment) Using Model Trained on Randomized Labels

**S132 CAMRand 26 14 DS+CHIR.** CAM Activation Maps of Colony 26 14 (DS+CHIR Treatment) Using Model Trained on Randomized Labels

**S133 CAMRand 26 15 DS+CHIR.** CAM Activation Maps of Colony 26 15 (DS+CHIR Treatment) Using Model Trained on Randomized Labels

**S134 CAMRand 26 16 DS.** CAM Activation Maps of Colony 26 16 (DS Treatment) Using Model Trained on Randomized Labels

**S135 CAMRand 26 17 DS+CHIR.** CAM Activation Maps of Colony 26 17 (DS+CHIR Treatment) Using Model Trained on Randomized Labels

**S136 CAMRand 26 18 DS.** CAM Activation Maps of Colony 26 18 (DS Treatment) Using Model Trained on Randomized Labels

**S137 CAMRand 26 19 DS+CHIR.** CAM Activation Maps of Colony 26 19 (DS+CHIR Treatment) Using Model Trained on Randomized Labels

**S138 CAMRand 26 20 DS+CHIR.** CAM Activation Maps of Colony 26 20 (DS+CHIR Treatment) Using Model Trained on Randomized Labels

**S139 CAMRand 26 21 DS+CHIR.** CAM Activation Maps of Colony 26 21 (DS+CHIR Treatment) Using Model Trained on Randomized Labels

**S140 CAMRand 26 22 DS+CHIR.** CAM Activation Maps of Colony 26 22 (DS+CHIR Treatment) Using Model Trained on Randomized Labels

**S141 CAMRand 26 23 DS+CHIR.** CAM Activation Maps of Colony 26 23 (DS+CHIR Treatment) Using Model Trained on Randomized Labels

**S142 CAMRand 26 24 DS.** CAM Activation Maps of Colony 26 24 (DS Treatment) Using Model Trained on Randomized Labels

**S143 CAMRand 26 25 DS.** CAM Activation Maps of Colony 26 25 (DS Treatment) Using Model Trained on Randomized Labels

**S144 CAMRand 26 26 DS.** CAM Activation Maps of Colony 26 26 (DS Treatment) Using Model Trained on Randomized Labels

**S145 CAMRand 26 27 DS+CHIR.** CAM Activation Maps of Colony 26 27 (DS+CHIR Treatment) Using Model Trained on Randomized Labels

**S146 CAMRand 26 28 DS.** CAM Activation Maps of Colony 26 28 (DS Treatment) Using Model Trained on Randomized Labels

**S147 CAMRand 26 29 DS+CHIR.** CAM Activation Maps of Colony 26 29 (DS+CHIR Treatment) Using Model Trained on Randomized Labels

**S148 CAMRand 26 30 DS.** CAM Activation Maps of Colony 26 30 (DS Treatment) Using Model Trained on Randomized Labels

**S149 CAMRand 26 31 DS+CHIR.** CAM Activation Maps of Colony 26 31 (DS+CHIR Treatment) Using Model Trained on Randomized Labels

**S150 CAMRand 26 32 DS.** CAM Activation Maps of Colony 26 32 (DS Treatment) Using Model Trained on Randomized Labels

**S151 CAMRand 26 33 DS+CHIR.** CAM Activation Maps of Colony 26 33 (DS+CHIR Treatment) Using Model Trained on Randomized Labels

**S152 CAMRand 26 34 DS.** CAM Activation Maps of Colony 26 34 (DS Treatment) Using Model Trained on Randomized Labels

